# Deep Phenotyping: Deep Learning for Temporal Phenotype/Genotype Classification

**DOI:** 10.1101/134205

**Authors:** Sarah Taghavi Namin, Mohammad Esmaeilzadeh, Mohammad Najafi, Tim B. Brown, Justin O. Borevitz

## Abstract

High resolution and high throughput, genotype to phenotype studies in plants are underway to accelerate breeding of climate ready crops. Complex developmental phenotypes are observed by imaging a variety of accessions in different environment conditions, however extracting the genetically heritable traits is challenging. In the recent years, deep learning techniques and in particular Convolutional Neural Networks (CNNs), Recurrent Neural Networks (RNNs) and Long-Short Term Memories (LSTMs), have shown great success in visual data recognition, classification, and sequence learning tasks. In this paper, we proposed a CNN-LSTM framework for plant classification of various genotypes. Here, we exploit the power of deep CNNs for joint feature and classifier learning, within an automatic phenotyping scheme for genotype classification. Further, plant growth variation over time is also important in phenotyping their dynamic behavior. This was fed into the deep learning framework using LSTMs to model these temporal cues for different plant accessions. We generated a replicated dataset of four accessions of Arabidopsis and carried out automated phenotyping experiments. The results provide evidence of the benefits of our approach over using traditional hand-crafted image analysis features and other genotype classification frameworks. We also demonstrate that temporal information further improves the performance of the phenotype classification system.

## 1 Introduction

Plant productivity must increase dramatically this century, while using resources more efficiently, to accommodate the ever-growing demand of a more affluent and growing human population. Precision breeding, via selecting advantageous genomic variants, will help improve plant productivity and efficiency but it relies on a detailed understanding of the genotype to phenotype relationship [1]. Here, automated phenotyping in high resolution across large mapping populations in multiple environments is needed to determine the underlying trait x genotype relationship. We have developed *climate chambers*, which maintain diurnal and seasonal climate signals but remove the weather noise plaguing field studies. These chambers have automated image capture capability to constantly monitor plants throughout their entire life cycle [2].

*Arabidopsis thaliana* is one of the model organisms used for studying plant biology, and it now has genomes sequences from 1000s of accessions [3]. Since the growth patterns of this plant are easily observable (especially from top-view), it is a very useful model for automated phenotyping. Previous work on *phenotyping* different *genotypes* (*accession*) have mostly used biologist specified, ’hand-crafted’ image features such as number of leaves, leaf area, compactness, roundness, etc. [4, 5, 6, 7, 8]. These features are computed either manually or via custom image processing algorithms. Their output may then be passed to a classifier. The main weakness of using hand-crafted descriptors is that although they are readily interpretable, they may be missing or incorrectly measuring the actual features that are variable among accessions. Furthermore, the custom image processing methods to extract the hand crafted features may not work as well when run on other experiments and may be difficult to generalize to more heterogeneous data sets [9].

Problems with hand crafted features have been addressed in the past few years by harnessing the power of deep learning *Convolutional Neural Networks* (CNNs) in particular [10, 11, 12, 13, 14], although difficulties with interpretation of the machine learned traits and over-fitting to a particular experiment remain. CNNs automatically find and extract the most descriptive features from the data during the training process. In other words, both feature extraction and training steps are performed simultaneously and hence, the system tries to find the features that minimize the loss criterion of the phenotyping problem. As a result, novel features for accession recognition are revealed in this process. However, in order for a machine to learn a good set of features, a very large training dataset is required.

CNNs are great for classification and segmentation of images, but they are unable to properly model dynamic systems, such as time lapse video in our case. Although CNNs can not encode temporal dependency of successive image frames, this problem can be addressed by using a *Recurrent Neural Network* (RNN) in which, each image frame is processed and analyzed by a neural cell and the information of each cell is circulated to the succeeding cells. RNNs, and in particular *Long Short-Term Memorys* (LSTMs, which are explained in detail in Sec. 3.2.1) have demonstrated potential in computer vision for analysis of dynamic systems [15, 16, 17, 18, 19]. In this study we utilize LSTMs to carefully model the growth patterns of plants.

## 2 Background

Research has focused on automatic plant phenotyping and classification using high-throughput systems. Classification of growth phenotypes based on data from known planted genotypes represents a typical experimental design where the aim is to obtain measures that maximize signal between genotypes relative to environmental error within biological replicates of the same genotype. Advanced image processing using machine learning techniques have become very popular in phenotyping qualitative states [20, 21, 22, 23, 24] while there are still many prospective needs and goals [25, 26, 27, 28, 29] to be experimentally explored in plants. A number of recent studies have presented high-throughput systems for plant phenotyping [30, 31, 32, 33, 2] and also plant/leaf segmentation and feature extraction [34, 35, 36, 37].

Plant classification has attracted researchers from the computer vision community [38, 39, 40, 41] given its importance in agriculture and ecological conservation. There are several studies of plant classification built on the pictures of individual plant leaves [42, 43, 44, 45]. Approaches to recognize plant disease [46, 47], symptoms of environmental stress [31, 48], and differentiation of crops from weeds [49, 50] have been previously studied. Normally three primary steps of plant/leaf segmentation, feature extraction, and classification are involved in these studies. The performance of the whole phenotyping pipeline depends on the performance and interaction among each of the three elements.

In the past few years, deep learning methods and in particular, Convolutional Neural Networks have achieved state-of-the-art results in various classification problems, and has motivated scientists to use them for plant classification tasks as well [51, 52, 53, 54]). CNNs are able to learn highly discriminative features during the training process and classify plants, without any need for segmentation or hand-crafted feature extraction. In particular, [54] used a CNN for root and shoot feature identification and localization. The authors in [52] proposed Deep Plant framework which employs CNNs to learn feature representation for 44 different plant species using the leaves. However, all the above-mentioned studies in plant phenotyping, feature extraction and classification are all based on individual static images of the plants of different species. In other words, temporal information, such as the growth patterns, one of the key distinguishing factors between varieties within plant species, has not been previously taken into account. Temporal cues can be very helpful, especially for distinguishing between different plants that have similar appearances, e.g. for separating different accessions of a particular plant, which is often a very challenging task.

In order to account for temporal information, various probabilistic and computational models (e.g. Hidden Markov Models (HMMs) [55, 56, 57], rank pooling [58, 59, 60], Conditional Random Fields (CRFs) [61, 62, 63] and RNNs [64, 65, 66, 67]) have been used for a number of applications involving sequence learning and processing.

RNNs (and LSTMs in particular) are able to grasp and learn long-range and complex dynamics and have recently become very popular for the task of activity recognition. More specifically, [15, 16, 17, 18, 19] used LSTM in conjunction with CNN for action and activity recognition were shown to provide a significant improvement in performance over previous studies of video data. In this paper, we treat the growth and development of plants as an action recognition problem, and use CNN for extracting discriminative features, and LSTM for encoding the growth behavior of the plants.

## 3 Preliminary

In this section, we explain the fundamentals of deep structures used in this paper, including CNN, RNN and LSTM.

### 3.1 CNN

Figure 1 depicts the schematic of a Convolutional Neural network (Alexnet [68]). Each layer in this network consists of a set of parameters and neuronal activations. The parameters could be trainable or fixed, depending on the layer. Activations could be non-linear or linear, as described in more detail below. The CNN structure takes a tensor of three-dimensional data as its input, passes it through multiple sets of layers and then outputs a score that represents the semantic class label of the input data. For instance in a simple cat vs. dog classification task, the input could be the image of a kitty and the correct output would be a high score for the cat class.

**Figure 1:**
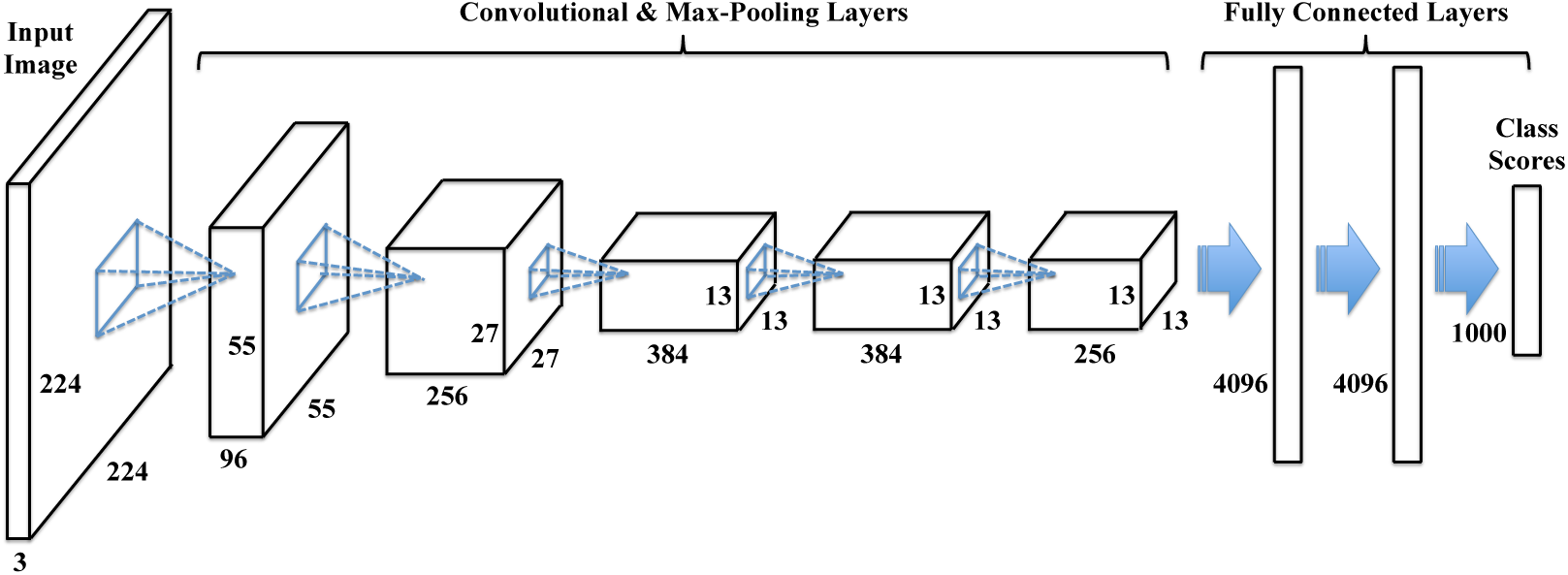
The schematic of Alexnet. A CNN often consists of convolutional layers, max-pooling layers and fully connected layers. The output of each convolutional layer is a block of 2D images (a.k.a. feature maps), which are computed by convolving previous feature maps with a small filter. The filter parameters are learned during the training process. The last few layers of CNN are densely connected to each other, and the class scores are obtained from the final layer.

In our application, we feed the CNN with top-view images (with three color channels) from plants. Next we introduce the main layers of a CNN.

#### 3.1.1 Convolutional Layer

This layer is computed by applying multiple filters to the input image, i.e. sliding the filter window over the entire input image. Different filters can have different parameters, which lets them detect and learn different image features. For example, one filter could be in charge of spotting vertical edges, while another one could detect horizontal edges. The output of this layer is called a feature map.

Filters are normally designed to be small (3 × 3, 5 × 5, 7 × 7, …), to reduce the number of parameters in the system. As a result, regardless of the size of the input image, the parameter size remains limited. This is in contrast to the design of a fully connected neural network where all the units in the previous layer are connected to every unit in the next layer with unique parameters, which leads to a sizable parameter set.

#### 3.1.2 Max Pooling Layer

Each feature map obtained from the convolutional layer, is an indicator of a particular feature in different locations of the input image. We normally want our descriptors to be robust against minor displacements of the input data. This is addressed by adding a *max pooling layer* to the network, which downsamples the feature maps. In other words, it reduces small patches of the feature map into single pixels. If a feature is detected anywhere within the patch, the downsampled patch fires a detection of that feature (local invariance).

A more practical benefit of the pooling layer is that, reducing the size of the feature maps leads to a significant decrease in the number of parameters, which in turn controls overfitting and also speeds up the training process. Another advantage of pooling layer is that it helps the network to detect more meaningful and high-level features as it moves on to the deeper layers. In this structure, the first layer has detected low level features like edges, whereas the next layer could grab more sophisticated descriptors like leaves or petiole, and the layer after has learned high-level features that are able to describe the whole plant.

#### 3.1.3 Fully Connected Layer

After a sequence of multiple convolution and pooling layers, the size of input data is shrunk dramatically which is suitable as input to a fully connected (dense) layer. The resulting feature maps up to this point of the network are vectorized and feed a multi-layer fully connected neural network, whose last layer (a.k.a classification layer or softmax layer) denotes the scores of the class labels in our problem.

The last fully connected layer is in charge of computing the scores for each class label. Each neuron in this layer represents a category in the classification problem, and its class probability can be computed by applying a *softmax* function to its inputs from the previous layer.

#### 3.1.4 CNN structure

The structure of a CNN (number of different layers, size of the filters, size of the fully connected layers, etc.) may vary depending on the application and the size of the training data. During the past few years, several architectures have been proposed and shown to work quite well for image classification and segmentation problems, among which AlexNet [68], VggNet [69] and ResNet [70] are the most notable ones.

Figure 1 shows the schematic of AlexNet, which has five convolution layers, three of which are followed by max pooling layers. It also features three fully connected layers. This is the network that first attracted the attention of researchers to the potential of CNNs, by winning the ImageNet Large Scale Visual Recognition Competition (ILSVRC) by a big margin [71], compared to the models with hand-crafted features.

### 3.2 RNN

Figure 2 illustrates a simple RNN that models a temporal data with three time points. In this representation, each time step is portrayed by a block of neurons, which receives two inputs respectively from the observed frame at that time, and the temporal cues propagated from previous times points. A fully connected neural network is embedded within each RNN cell to analyze the visual information of each frame together with the information that is received from previous times, to obtain the system state at each time frame. Let **x**(*t*), **h**(*t*) and **y**(*t*) denote the visual input data, the output of RNN cell and the class label of the sequential data, respectively, at time *t*. Then the RNN can be expressed as

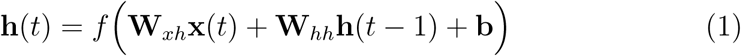

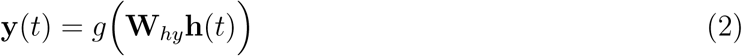

where **W**_*xh*_, **W**_*hh*_ and **W**_*hy*_ are the neural network parameters, **b** is a bias vector, and *f* and *g* are element-wise non-linear functions which are often set to *hyperbolic tangent* (*ϕ*) and *sigmoid* (*σ*), respectively.

**Figure 2:**
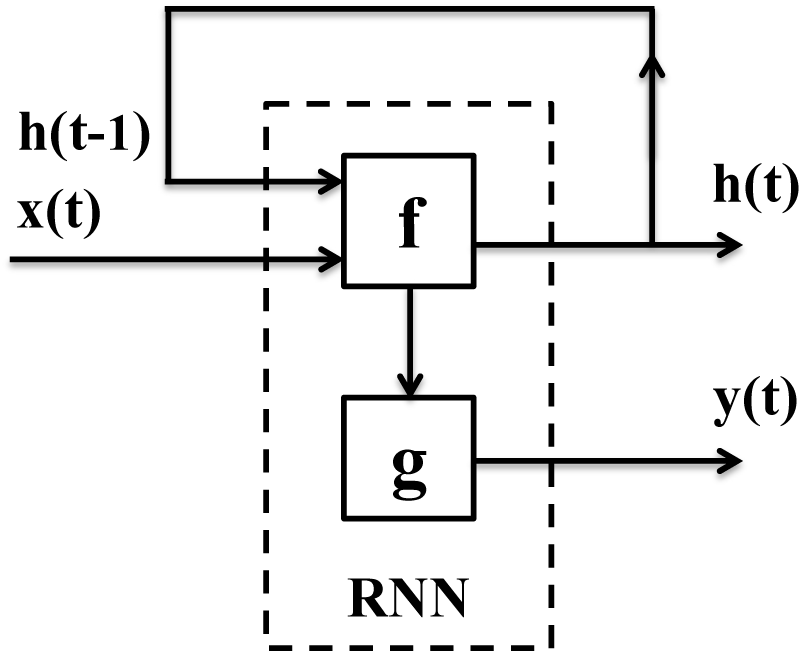
The structure of an RNN. The system at each time-point is updated based on the current input data and the status of the system at the previous time-point. Here, *f* and *g* are element-wise non-linear functions which are often set to *hyperbolic tangent* (*ϕ*) and *sigmoid* (*σ*), respectively.

What makes this structure more interesting is that we can readily integrate RNN with a CNN, by feeding the visual input of the RNN cell with the pre-trained CNN features of the image frame at that time point.

#### 3.2.1 LSTM

The main shortcoming of standard RNNs (Figure 2) is that they can not encode temporal dependencies that prolong to more than a limited number of time steps [72]. In order to address this problem, a more sophisticated RNN cell named Long Short-Term Memory (LSTM) has been proposed to preserve the useful temporal information for an extended period of time.

An LSTM, as depicted in Figure 3, is equipped with a memory cell and a number of gates. The gates control when a new piece of information should be written to the memory or how much of the memory content should be erased. Similar to a standard RNN, the state of the system at each time point is computed by analyzing the visual input at that time point, together with the output of previous cell and also the content of the LSTM memory, which is referred to as **c**(*t*). Given **x**(*t*), **h**(*t*) and **c**(*t*), the LSTM updates are defined as

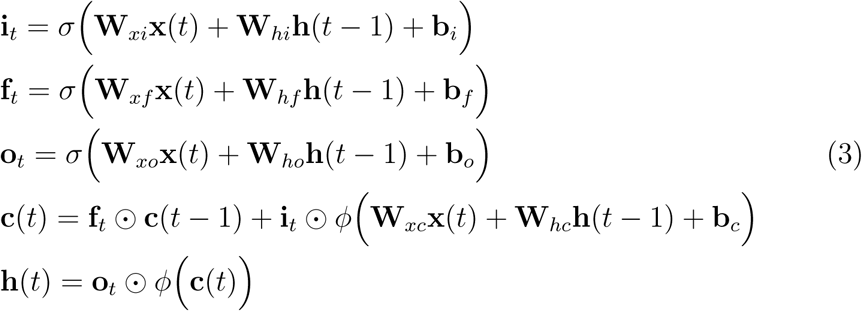

**Figure 3:**
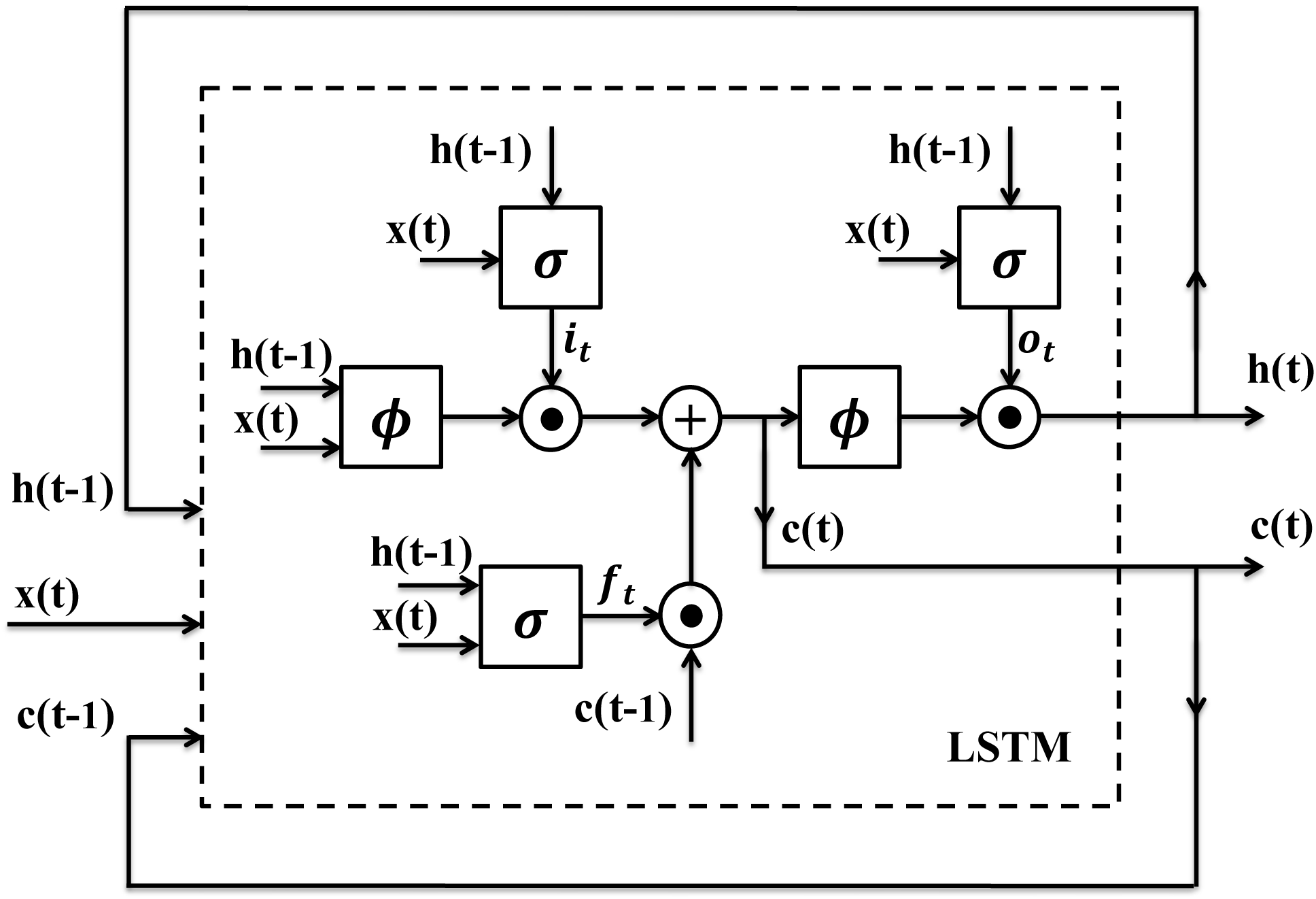
The structure of an LSTM. The system at each time-point is updated based on the current input data, the status of the system at the previous time-point, and the content of the memory. Here, *ϕ* and *σ* are *hyperbolic tangent* and *sigmoid* functions, respectively, and ⊙ stands for the element-wise multiplication.

In these equations, **i**_*t*_, **f**_*t*_ and **o**_*t*_ denote input gate, forget gate and output gate respectively. The input gate controls how much of the new input data should be recorded into the memory, whereas the forget gate decides how much of the old memory should be preserved at each time. The output of the LSTM cell is also computed by applying the output gate to the memory content. This sophisticated structure enables LSTM to perceive and learn long-term temporal dependencies. Note that ⊙ in Eq. 3 indicates an element-wise multiplication.

After seeing a sufficient number of data sequences in the training phase, LSTM learns when to update the memory with new information or when to erase it, fully or partially. LSTMs can model various sequential data very easily, unlike other complicated and multi-step pipelines. Furthermore, they can be fine-tuned similar to CNNs. These benefits has made LSTMs very popular in the recent years for modelling data sequences. In this paper, we propose a CNN-LSTM structure (Figure 4) to build a plant classification system, which is explained in more detail in Sec. 4.1.

**Figure 4:**
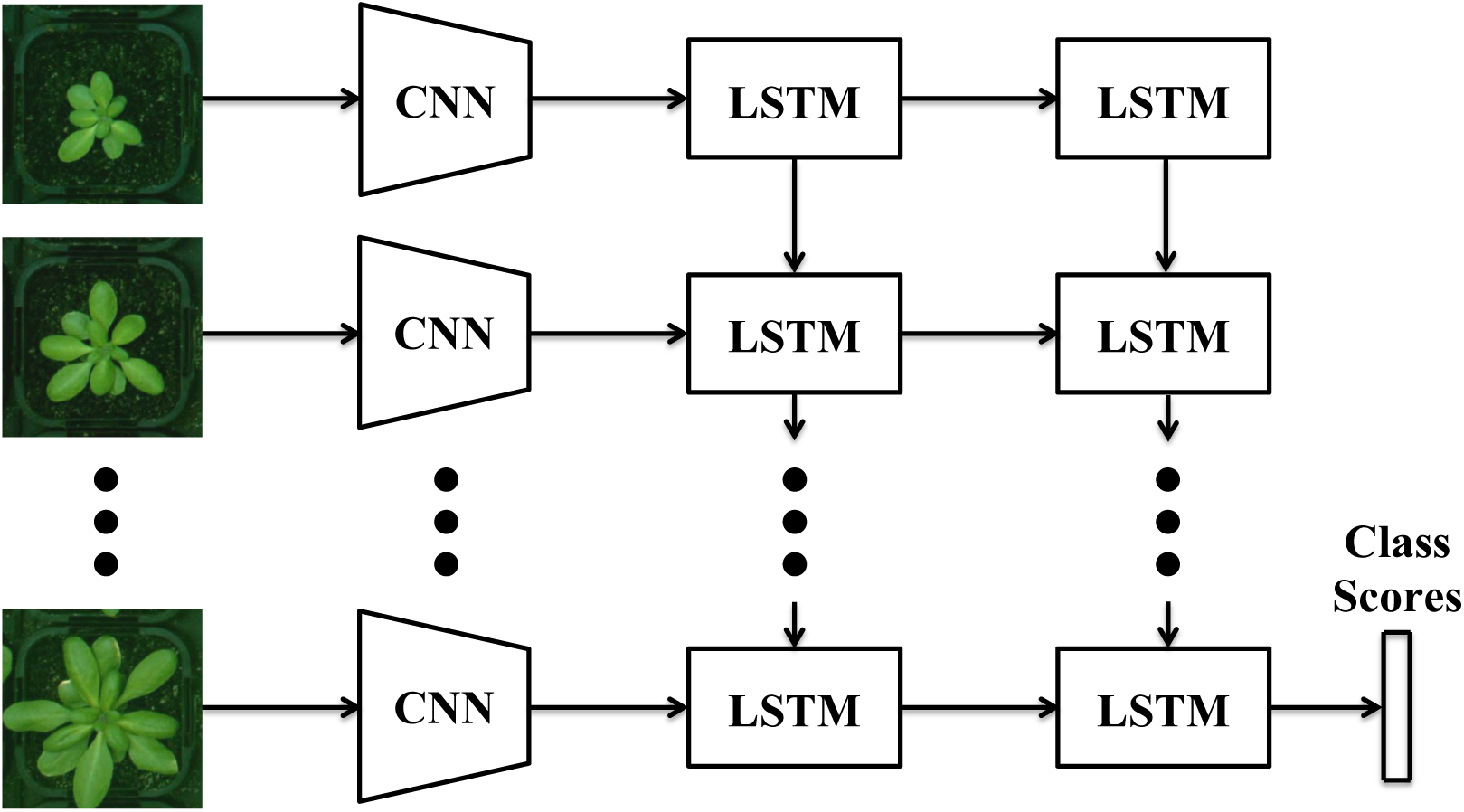
The CNN-LSTM structure. The CNNs extract deep features of the plant images and then, the growth pattern of the plant is modeled using LSTMs. Finally the genotype with highest class score is selected.

## 4 Our Approach

We aim to propose an automatic accession classification framework, using the deep visual features of the plants (which are trained specifically for the accession categories) as well as the temporal cues of the plant growth sequences. To this end, in this section we introduce the CNN-LSTM model and then explain how to train this model.

### 4.1 CNN-LSTM Network

In this section, we describe the proposed framework for genotype classification, which is composed of a deep visual descriptor (using a CNN), and an LSTM which can recognize and synthesize temporal dynamics in an image sequence such as growth rate, number of leaves, and texture changes. As depicted in Figure 4, our approach is to first pass each individual frame of the plant image sequence through the deep visual descriptor (CNN) to produce a fixed-length vector representation. This fixed-length vector embodies the features of each individual plant, which are extracted after fine-tuning step (as explained in Sec. 4.2). The outputs of CNN for the sequence of pot images are then passed onto a sequence learning module (LSTM). At this stage, the LSTM attempts to classify the plants via analyzing the sequences of the features that are extracted from image frames and by taking into account their temporal variations. In other words, the proposed CNN-LSTM structure encodes the activity of the plants during their growth period to model the relationships between their phenotypes and genotypes.

The proposed model can automatically classify plants into the desired categories, given only the plant images. Note that our approach can be easily extended to the cases, where more classes are involved, just by performing the training phase for the new set of classes. Extending the model to applications other than plant classification is just as easy, where one can simply modify the target layer of the network to fit that particular problem. This is counter to the conventional phenotyping methods, where one is required to find relevant hand-crafted features for each individual application.

### 4.2 CNN Training

The goal of training is to find the values of network parameters such that the predicted class labels for the input data are as close as possible to their ground truth class labels. This, however, is a very challenging task since CNNs normally have a vast number of parameters to be learned. AlexNet for instance is built on more than 60 millions parameters. Training a system with this many parameters requires a massive number of training images as well.

There are a few publicly available datasets that provide sufficient number of images for training CNN architectures, among which ImageNet-ILSVRC is very popular. It is a subset of much larger ImageNet dataset and has about 1.2 millions images selected from 1000 different categories. However, in many problems we do not have access to a large dataset, and this prevents us from properly training a CNN for them.

It is shown if we initialize the network using the parameters of a pre-trained CNN (a CNN that is already trained on a big dataset like ImageNet), and then train it using the limited dataset in our problem, we can achieve very good performance. In particular, we can rely on the basic features that the CNN has learned in the first few layers of the network on ImageNet, and try to re-train the parameters in the last few layers (normally fully connected layers) such that the network could be fit to our specific problem. This method is often referred to as *fine-tunning*, which speeds up the training process and also prevents overfitting of the network to a relatively small dataset.

Note that in many image classification problems, it is very common to preserve all the layers and parameters of a pre-trained CNN, and only replace the last layer that represents the 1000 class labels of ImageNet with the class labels in our specific problem. Then only the parameters of the classification layer are learned in the training phase, and the rest of the parameters of the network are kept fixed to the pre-trained settings. In fact here we assume that the deep features that are previously learned on ImageNet dataset can describe our specific dataset quite well, which is often an accurate assumption. The outputs of the layer before the classification layer of a CNN are sometimes refereed to as *pre-trained CNN features*.

In this work, we chose to fine-tune a pre-trained CNN using the top-view images of the plants, in order to learn more discriminant features for distinguishing different accessions.

#### Data Augmentation

When a dataset has a limited number of images, which is not sufficient for properly training the CNN, it makes the network vulnerable to overfitting. In order to synthetically increase the size of the training data, we can use a simple and common technique, called *Data Augmentation*. In this procedure, we rotate each image in the dataset by 90°, 180° and 270° around its center and add it to the dataset.

### 4.3 Deep Feature Extraction

Our goal is to classify plants into different genotypes (Accessions), as depicted in Figure 5. First, we need to train a CNN on our plant dataset to learn the deep features that are fed to the LSTM cells. We use Alexnet, which is pre-trained on ImageNet to provide us with very descriptive features. Note that we choose AlexNet over deeper network such as VggNet or ResNet, because it has fewer parameters to learn, which better suits our limited dataset. We then replace the last layer of AlexNet with a layer of *L* neurons to adapt the network to our application.

**Figure 5:**
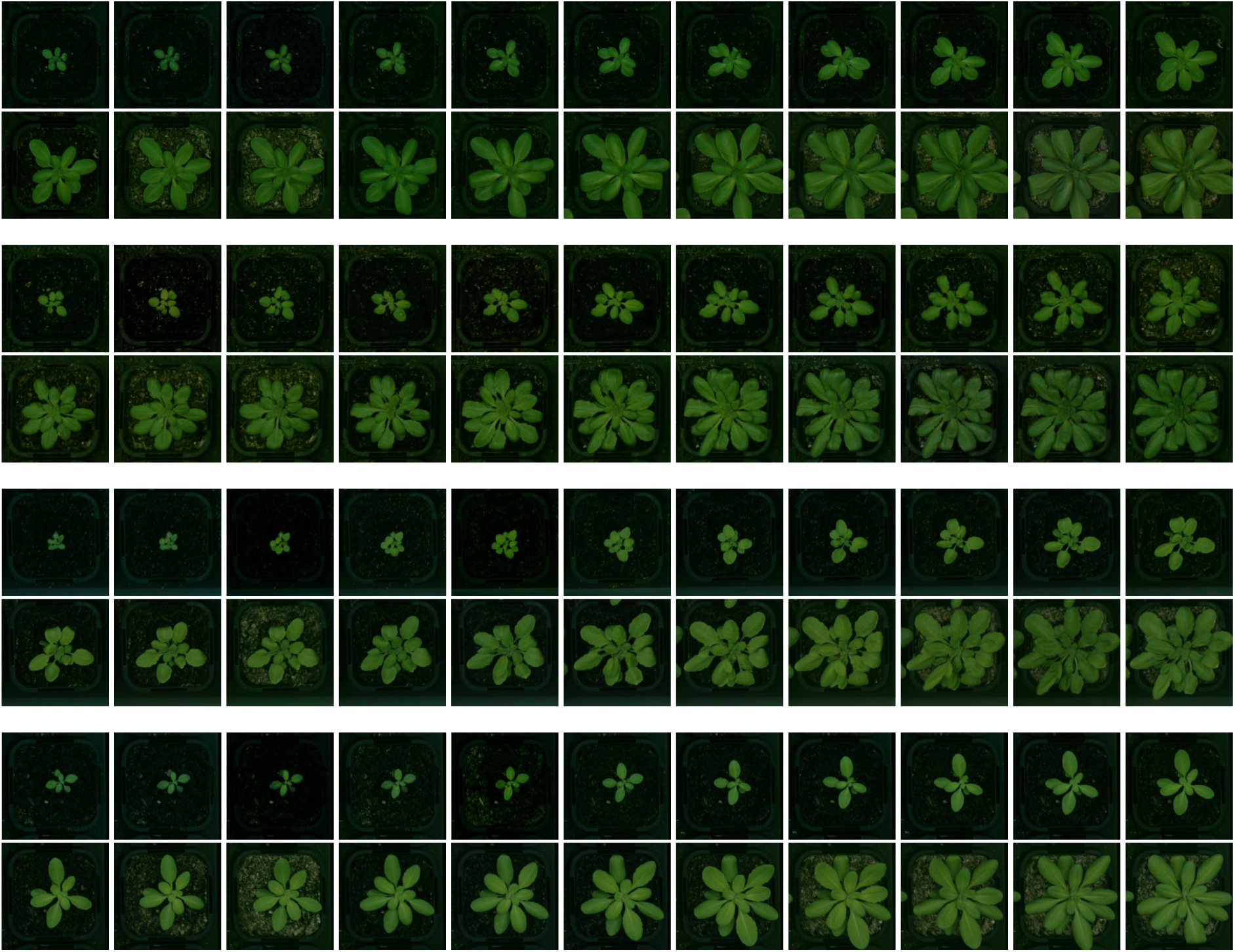
Samples of sequence data from various accessions. Examples of sequence data including 22 successive top-view images of 4 different categories of Arabidopsis thaliana. Successive images are recorded at 12:00pm of every day. From top to bottom, accessions are: SF-2, CVI, Landsberg(Ler), and Columbia (Col).

Our dataset is composed of sequences of images captured from the plants in different days while they grow. We initially break down image sequences of the plants into individual images in order to build a CNN training dataset, and then use data augmentation to extend the size of this dataset, as explained in Sec. 4.2. However, since plants change in size a lot during their growth, the decomposed images from the plant sequences are not sufficiently consistent to form a proper training dataset for a genotype. This makes CNN training very difficult, if not impossible, particularly in our case where the total size of the training set is very limited.

We account for this intra class variability by splitting each genotype class into a class set of that genotype in multiple area sizes. The area is computed by counting the total number of pixels that belong to the plant, and is computed by segmenting the image. The segmentation outputs for a sample plant sequence is shown in Figure 7. Plant segmentation process is explained in Sec. 5.2. Another factor that could have been considered for breaking down each genotype into smaller and more consistent categories, is the day when the plant is observed and its image is captured. This factor, which somehow encodes the growth rate of the plant, is not however purely dependent of the genotypes and is heavily affected by environment conditions such as germination occurring on different days. Note that even though the experiments are conducted inside growth chambers where environment conditions are to be controlled, the plants still show variability.

**Figure 6:**
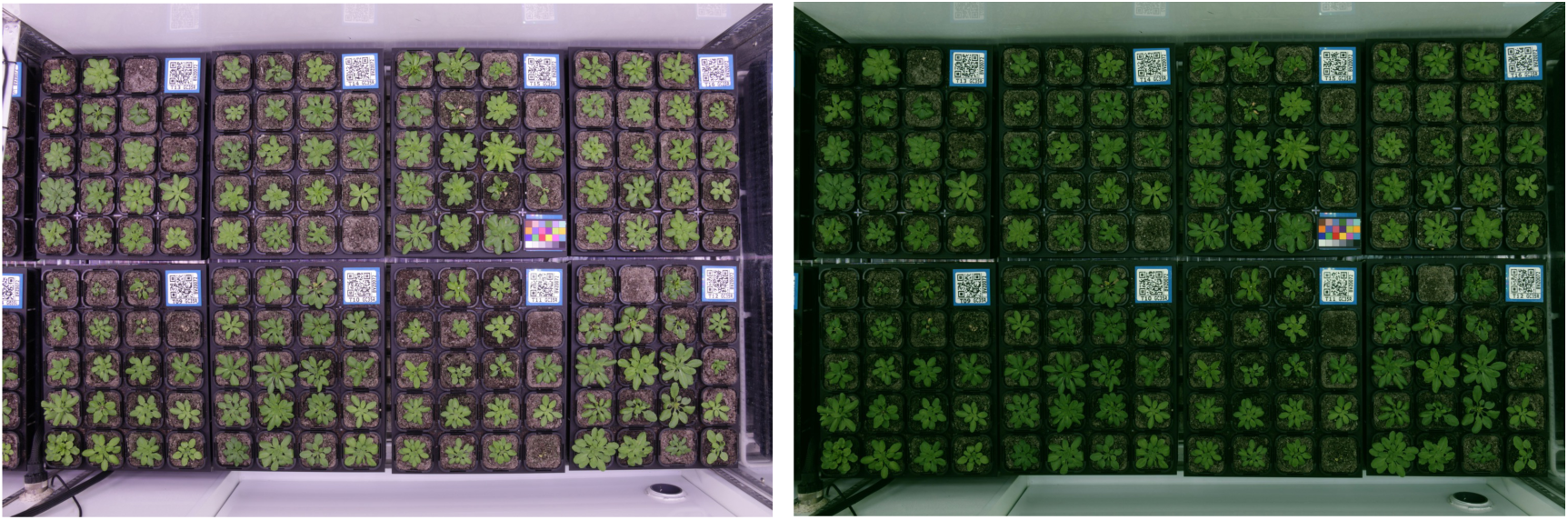
Growth chamber. Left: The original picture of a growth chamber; Right: The result of camera distortion removal and color correction step.

**Figure 7:**
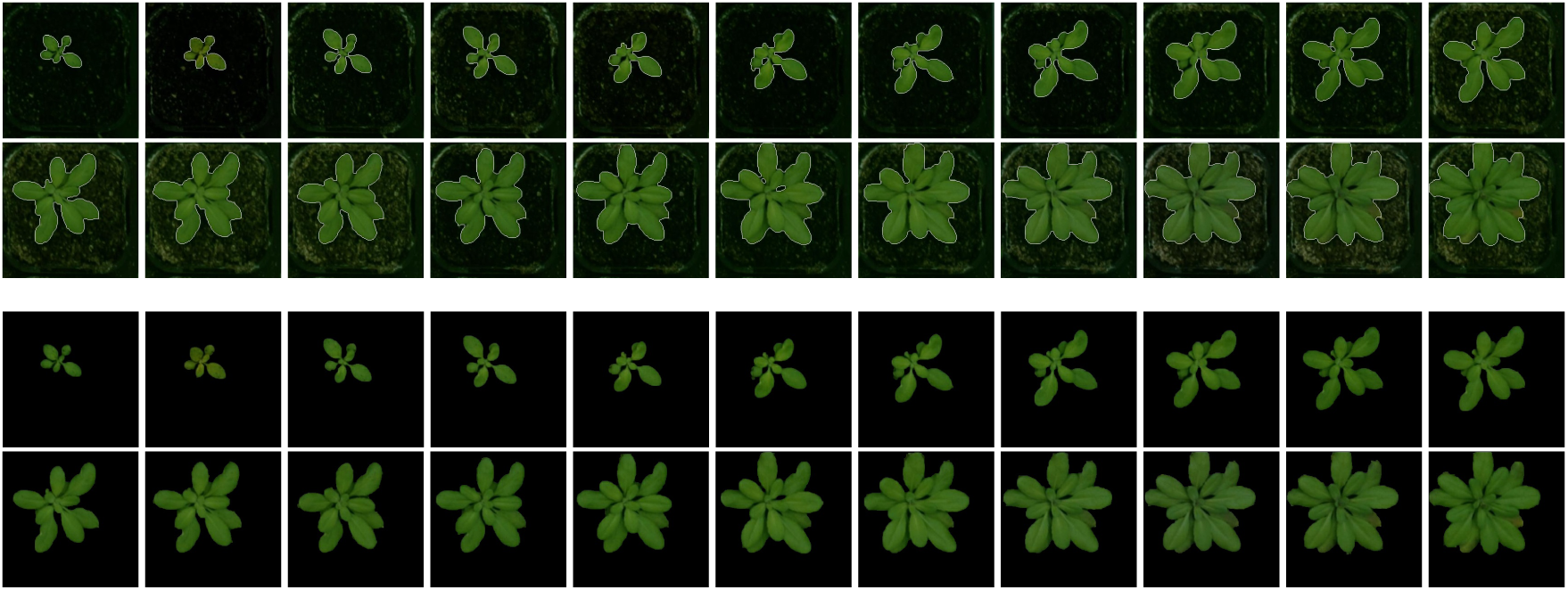
Plant Segmentation. The result of segmentation step is shown in this figure; top: plant contours, bottom: plant segments.

Given the area as a proper class divider, each genotype category is split into five sub-classes based on the plant areas, which means the CNN training is performed on *L*×5 classes. Once the CNN is trained, for each plant image we can use the output of the last fully connected layer before the classification layer, as deep features of the plant and feed them into the corresponding time point of the LSTM, in our CNN-LSTM structure.

### 4.4 LSTM Training

In order to train the LSTM, we feed it with sequences of deep features that are computed by applying the approach in Sec. 4.3 to the training image sequences. The system is then optimized to predict the true class label of the plants based on the information of the entire sequence. Note that we deepen the sequence learning module by adding another layer of LSTM to the structure (Figure 4). This enhances the ability of the proposed system to learn more sophisticated sequence patterns and in turn, improves the classification accuracy.

## 5 Experiments

In this section, we first introduce the dataset and then explain the preprocessing and plant segmentation steps. Next, we report the accession classification results using the proposed CNN-LSTM method. In order to evaluate this method more thoroughly, we extract a set of hand-crafted features and investigate their performance in the accession classification task, compared to our CNN-LSTM framework that uses deep features. Furthermore, we report the results of a variant of our approach where the LSTM is replaced by a CRF, to have a more thorough temporal analysis of the proposed model.

### 5.1 Our Dataset

We generated a plant dataset which is comprised of successive top-view images of *L* = 4 different accessions of Arabidopsis thaliana, which are SF-2, CVI, Landsberg (Ler) and Columbia (Col), as depicted in Figure 5. An example growth chamber that is used in our experiments is depicted in Figure 6-a, which contains a color card for color correction, and each tray in the chamber is accompanied with a QR code. Every pot is constantly monitored via a Canon EOS 650D, which is installed above the chamber.

In this work, we use the pot images that are recorded at 12:00pm of every day to build the data sequence of each plant. We do not include more than one image per day, as it makes the sequences longer, and the classification process becomes more computationally expensive, while it does not add significant temporal information. The obtained sequence for each plant involves 22 successive top-view images.

A number of pre-processing steps are applied to the captured images before moving on to the classification task. The first step is *camera distortion removal* to eliminate image distortions, flattening the image so pots are equal sizes. Then the images undergo a *color correction* process using the included color cards in the chambers. This step transforms the plant colors to make them appear as similar as possible to the real colors (Figure 6-b). Moreover, we use a *temporal matching* approach to detect trays and individual pots inside the trays, in order to extract the images of each pot and in turn generate the image sequence of the growing of each plant.

### 5.2 Plant Segmentation

For segmenting the plants we use the GrabCut algorithm [73], which is a method of distinguishing foreground from background based on the graph cuts [74]. In this algorithm, in addition to the input image, a bounding box that encompasses the foreground object should also be given as an input. Furthermore, a mask image with four intensity levels, representing definite background (0), definite foreground (1), probable background (2) and probable foreground (3) can also be provided as an auxiliary input to improve the segmentation.

Since the plants can be anywhere in the pots, especially when they grow large, we choose the bounding box to be as large as the input image to ensure no part of plants is missed. To generate the mentioned quaternary mask, the following approach is proposed.

First, the image is transformed from RGB into L\**a*\**b* colour space, as the plants and background are better distinguishable in *a* and *b* channels. Then, for each of the *a* and *b* components, image binarization using Otsu’s method [75] is performed; the outcome is two binary masks that highlights candidate foreground and background points for each of the channels. To ensure no part of the plants is mistakenly assumed as definite background, especially the leaf borders that could be faded into the soil in the images, next we use morphological dilation to expand the mask and this is then added to the binary mask. This leaves us with two masks, each having three intensity levels, 0: definite background, 1:probable background/foreground and 2: foreground.

The two masks are then combined to form the ultimate mask using the mapping in Table 1. The obtained mask is then used in the GrabCut algorithm to segment the plants. Finally, morphological opening and closing operations are applied to remove unwanted holes and blobs. The segmentation results for a sample sequence is shown in Figure 7.

**Table 1:** Combining the two binary masks computed from *a* and *b* color channels to produce the final mask for Grab-cut segmentation algorithm.

|  | 0 | 1 | 2 |
| --- | --- | --- | --- |
| 0 | Definite Background | Probable Background | Probable Foreground |
| 1 | Probable Background | Probable Background | Probable Foreground |
| 2 | Probable Foreground | Probable Foreground | Definite Foreground |

### 5.3 CNN-LSTM

We implemented our deep structure using Theano [76] and Keras [77]. We trained the parameters of the CNN using Stochastic Gradient Descent (SGD) method in mini-batches of size 32 and with a fixed learning rate of 0.001, a momentum of 0.9, and a weight decay of 1e-6. Similarly, we used SGD for the training of LSTM and trained it in mini-batches of size 32 with a fixed learning rate of 0.01, a momentum of 0.9, and a weight decay of 0.005. The LSTM is equipped with 256 hidden neurons. Table 2 illustrates the results of using our CNN-LSTM structure for accession classification, compared to the case where only CNN is used for classification and temporal information is ignored. Adding the LSTM to our structure has led to a significant accuracy boost (76.8% to 93%), which demonstrates the impact of temporal cues in accession classification. Table 2 reports comparisons with other benchmarks, which are explained in more detail in the next sections.

**Table 2:** The performance of our deep phenotyping system (CNN + LSTM) compared to other baseline methods. Results are reported in percent(%).

|  | Sf-2 | Cvi | Ler-1 | Col-0 | Avg. |
| --- | --- | --- | --- | --- | --- |
| CNN | 71.0 | 86.0 | 74.4 | 76.0 | 76.8 |
| Hand-crafted Features + LSTM | 61.3 | 90.2 | 68.2 | 52.4 | 68.0 |
| Hand-crafted Features using SVM | 58.4 | 83.9 | 65.1 | 35.7 | 60.8 |
| CNN + LSTM | **89.6** | 93.8 | **94.2** | **94.2** | **93.0** |
| CNN + CRF | 84.3 | **96.1** | 90.4 | 89.4 | 87.6 |

### 5.4 Phenotyping using Hand-crafted Features

We conduct an experiment where hand-crafted features, which are extracted from the plant images, are fed to the LSTM instead of deep CNN features. Then we can evaluate the contribution of deep features in our framework.

The features, which are extracted from the segmented plant images, are as follows: Mean, Max and Min of RGB image; Mean of HSV image; area and perimeter of the plant; *roundness* of the plant which is the ratio between its area and perimeter; *compactness* which is the ratio between area and convex-hull area; *eccentricity* which is the ratio between the major axis and minor axis of the convex-hull; length of the ellipse with the same second moment as the region; and *extent* which is the ratio between the area and the bounding box.

Furthermore, we compute a set of Fourier descriptors [78] to describe the shapes of the leaves in terms of their contours. It is worth noting that we make the Fourier features invariant to translation by setting the centre element of the Fourier transform of the image contours to zero. In total, a vector of 1024 elements (composed of 512 real and 512 imaginary components of the Fourier transform) is extracted to represent the contour shape of each plant.

In addition, we employ a set of texture features using the Gray-Level Co-occurrence Matrix (GLCM) [79, 80]. These features are extracted from segmented image plants and as a result, the texture information of different accessions are taken into account in the classification process. The obtained features via this method are independent of gray-level scaling of images and therefore, invariant to various illuminations and lighting conditions [80, 81]. Each element of GLCM indicates the frequency of the adjacency of a particular pair of gray-level intensities. In this experiment, we considered adjacencies in four directions of 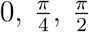 and 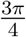, computed a GLCM for each direction, and then extracted three texture properties, *Energy*, *Contrast* and *Homogeneity* from each of the computed GLCMs. In total, this method provided us with 12 texture descriptors for each segmented plant.

The results of using hand-crafted features are reported in Table 2, which could be compared with the results of the proposed system (68% compared to 93%). Note that the quality of extracted hand-engineered features depends on how good the segmentation step is done. If the plants are not segmented properly, we may not obtain a reliable set of hand-crafted features, which in turn deteriorates the system performance even more.

The experimental results indicate the superiority of deep features compared to the above hand-engineered descriptors for accession classification. Note that we attempted to include a large array of various hand-crafted features in this experiment, but the classification system built on these descriptors was outperformed by our CNN-based classifier. Note that using a pure CNN-based classifier with no sequence learning module involved (no LSTM), led to a classification accuracy of 76.8%. This configuration outperforms the system with hand-crafted features (68%), and clearly indicates the benefit of using deep features over hand-engineered descriptors.

In addition, we perform another experiment where the temporal information of the plants are discarded and LSTMs are dropped from the structure. Then a Support Vector Machine classifier (SVM) is applied to the handcrafted features to predict the accession of each plant. This further degrades the classification performance of the system, as shown in Table 2.

### 5.5 CNN-CRF

The CRF is a popular probabilistic graphical model for encoding structural and temporal information of sequential data [82], and it has been widely used in the computer vision community [61, 62, 63, 83, 15, 84]. At its simplest form, this model encourages the adjacent elements in the spatial or temporal structure to take similar or compatible class labels and hence, it gives rise to a more consistent label for the whole structure (sequence).

In this work we studied the potential of the CRF for sequence analysis and compared it with RNNs in our sequence learning and accession classification experiment. To this aim, we fed the CRF with the previously computed deep features and reported its performance in the sequence classification task. Table 2 demonstrates the potential of CRFs for encoding the temporal dependencies in the sequential data, though they are still outperformed by our CNN-LSTM framework.

## 6 Conclusion

In this paper, we proposed a framework for automatic plant phenotyping based on deep visual features of the plants and also temporal cues of their growth patterns. Our experiments evidence the benefits of using deep features over hand-crafted features, and indicate the significance of temporal information in a plant classification task. Due to the generality of the proposed approach, it can be readily accommodated to other objectives and classification targets (based on genotypes or various environment conditions) and be trained on various datasets. Despite the deep learning demand for a large input dataset and our limited sequential data from different accessions, we presented a sophisticated deep network and an efficient method to train it. In the future, we plan to augment our dataset with more varying visual and sequential data to enhance the robustness of our system when dealing with more challenging classifications. We also intend to adapt our framework for a plant classification task based on growing environment conditions. Finally we plan to apply our trained classifier to a large set of accessions. The probability of each genotype state, SF-2, Cvi, Ler, Col, is a multivariate growth pattern phenotype of each accession, which can be decomposed into its causal genetic factors using Genome Wide Association Studies.

## 7 Acknowledgements

We thanks funding sources including, Australian Research Council (ARC) Centre of Excellence in Plant Energy Biology CE140100008, ARC Linkage Grant LP140100572 and the National Collaborative Research Infrastructure Scheme - Australian Plant Phenomics Facility.

